## Supplementary Material for "From complete cross-docking to partners identification and binding sites predictions"

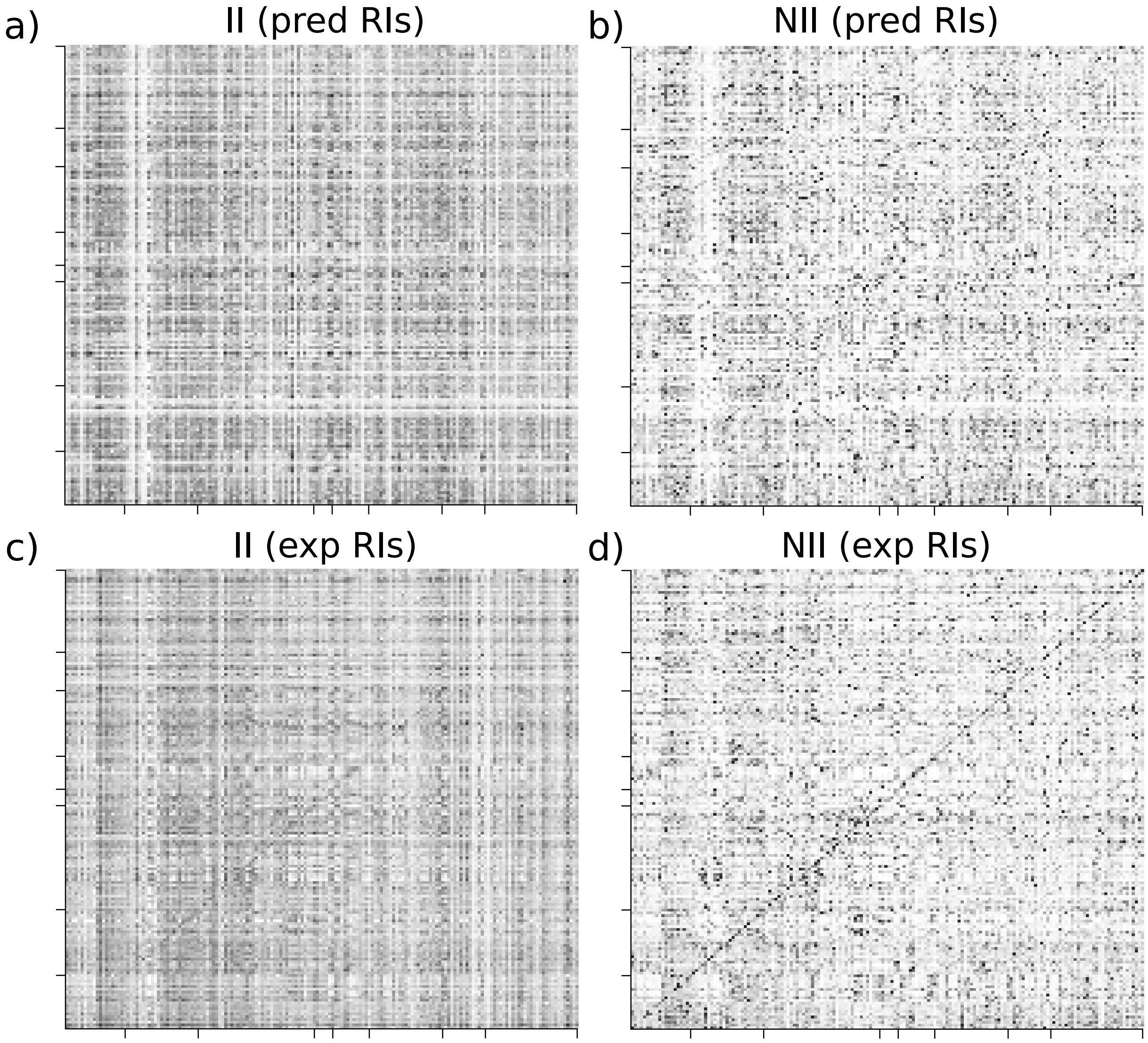


Figure S1: **Predicted interaction matrices for the PPDBv2.** **(a-b)** Matrices computed using predicted interfaces as references. **(c-d)** Matrices computed using experimental interfaces as references. The matrices on the left give interaction indices ($II$) and those on the right the normalized interaction indices ($NII$).

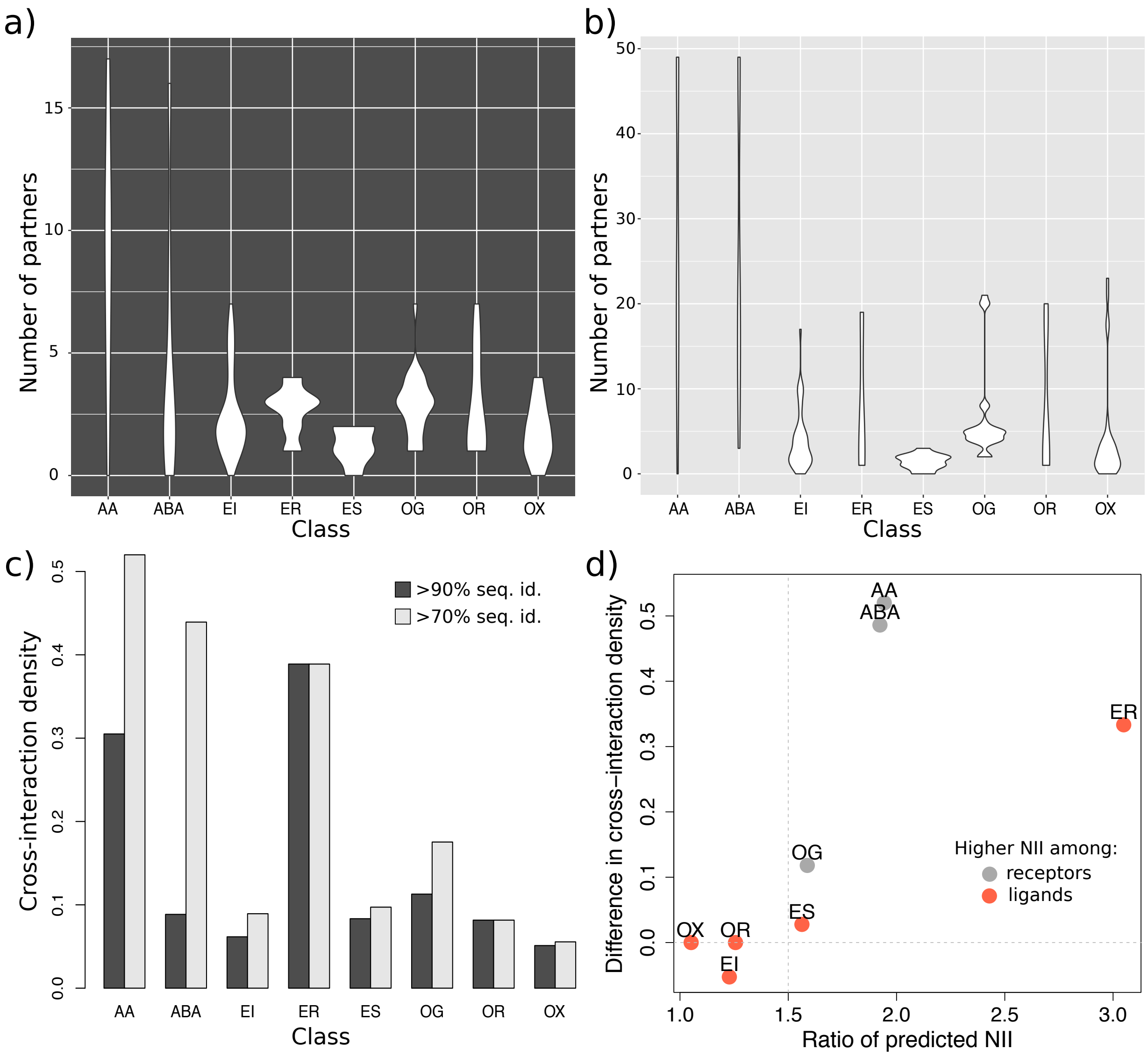


Figure S2: **Properties of the known interacting pairs**. **(a-b)** Distributions of the number of partners, for each protein within each subset, inferred by homology at 90% (a) and 70% (b) sequence identity levels. **(c)** Cross-interaction density, defined as the percentage of cells corresponding to a known interaction, within the matrix associated to each subset. The two grey tones indicate the sequence identity level. **(d)** Agreement between cross-interaction density and predicted NII values. In x-axis are reported the ratios $r_k = \max\left(\frac{\sum_{i,j\in S_k} NII_{R_i,R_j}}{\sum_{i,j\in S_k} NII_{L_i,L_j}}, \frac{\sum_{i,j\in S_k} NII_{L_i,L_j}}{\sum_{i,j\in S_k} NII_{R_i,R_j}}\right)$. For each subset $S_k$, $r_k$ reflects the difference in predicted interaction strengths among the receptors versus the ligands. When the dot is grey, it means the receptors are predicted to interact more with each other, while a red dot indicates that the ligands interact more. In y-axis are reported the different of cross-interactions densities between receptors and ligands, or reciprocally. When the value is positive, it means the tendency observed for the known interactions agrees with that observed for the predictions. For instance, antibodies are predicted to interact with each other twice more than antigens, and there are 50% more known interactions between them. Known interactions were determined with a sequence identity level of 70%.

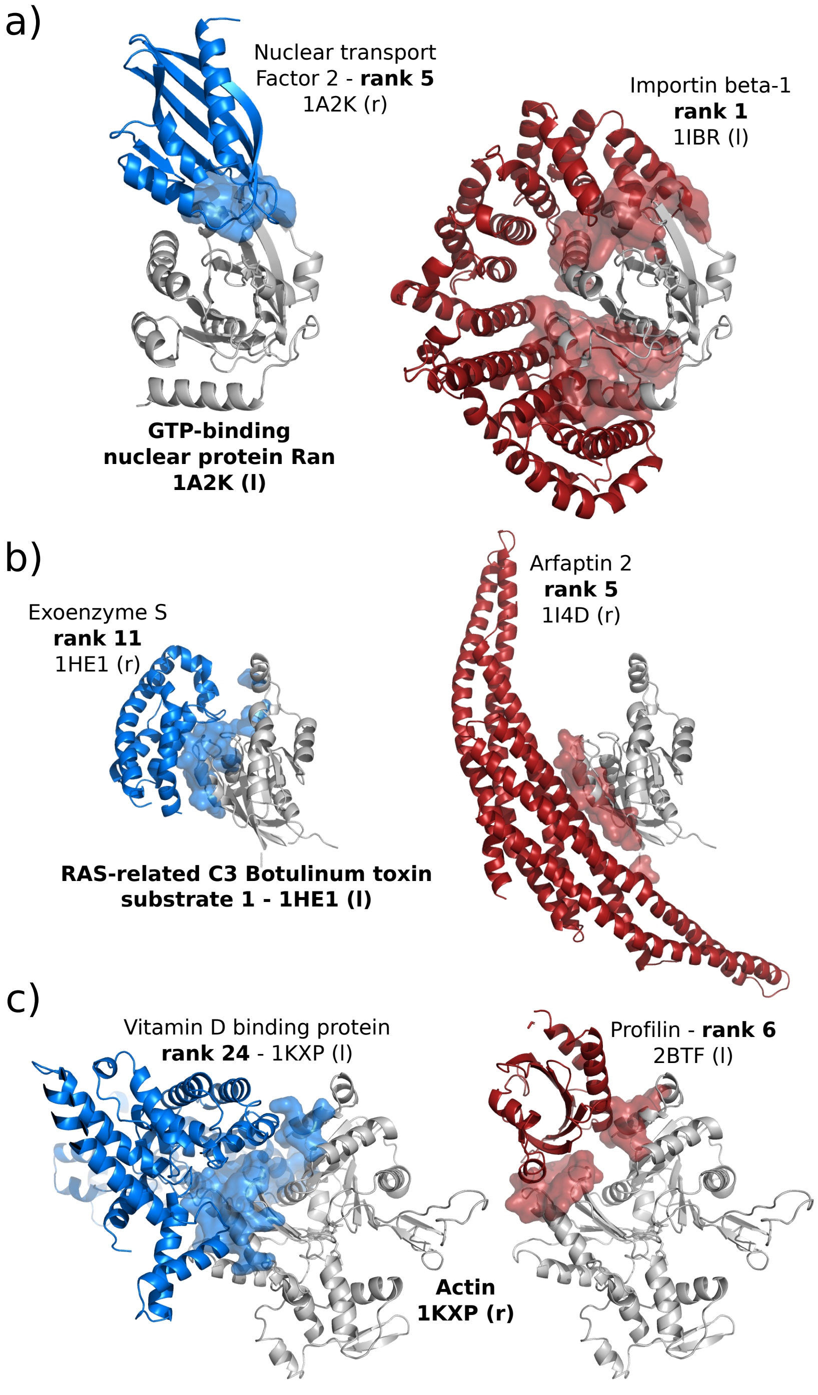


Figure S3: **Examples of annotated and homology-transferred interactions.** The query protein is represented as a grey cartoon. The cognate partner annotated in the PPDBv2 is shown in blue and a partner identified in the PDB by homology transfer (>90% sequence identity) is shown in dark red. In each case, the proteins come from the same functional class: **(a-b)** other-with-G protein, *OG*, **(c)** others, *OX*. The intra-class ranks of the partners are given.

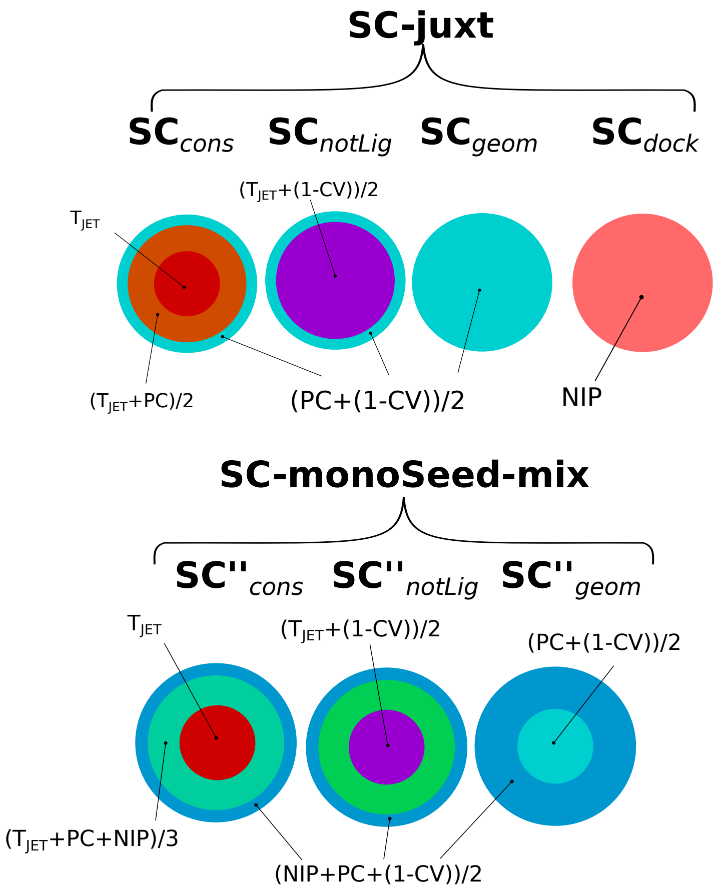


Figure S4: **Scoring schemes used to predict interfaces.** Each scoring scheme is depicted by a schematized representation of a predicted patch, where the different concentric layers correspond to different combinations of four residue-based descriptors. $T_{JET}$: evolutionary conservation. $PC$: physico-chemical properties. $CV$: circular variance. $NIP$: docking-inferred binding propensities. **Top left panel:** *SC-juxt* comprises four scoring schemes, three of them ($SC_{cons}$, $SC_{notLig}$ and $SC_{geom}$) using $T_{JET}$, $PC$ and $CV$ and the fourth one ($SC_{NIP}$) exclusively based on $NIP$. $SC_{cons}$ detects highly conserved seeds and extend them using physico-chemical properties and local geometry. $SC_{notLig}$ is a variant of $SC_{cons}$ including circular variance at the seed detection step to avoid buried ligand-binding pockets. $SC_{geom}$ disregards evolutionary conservation and detects protruding regions with good phyisco-chemical properties. All other scoring schemes are variants of $SC_{cons}$, $SC_{notLig}$ and $SC_{geom}$ including $NIP$ in different ways. **Top right panel:** *SC-mix* combines $NIP$ with the other descriptors at each step. **Bottom left panel:** *SC-monoSeed-mix* disregards $NIP$ to detect the seeds and then combines it with the other descriptors. **Bottom right panel:** *SC-dockSeed-mix* relies exclusively on $NIP$ to detect seeds and then uses a combination of the four descriptors.

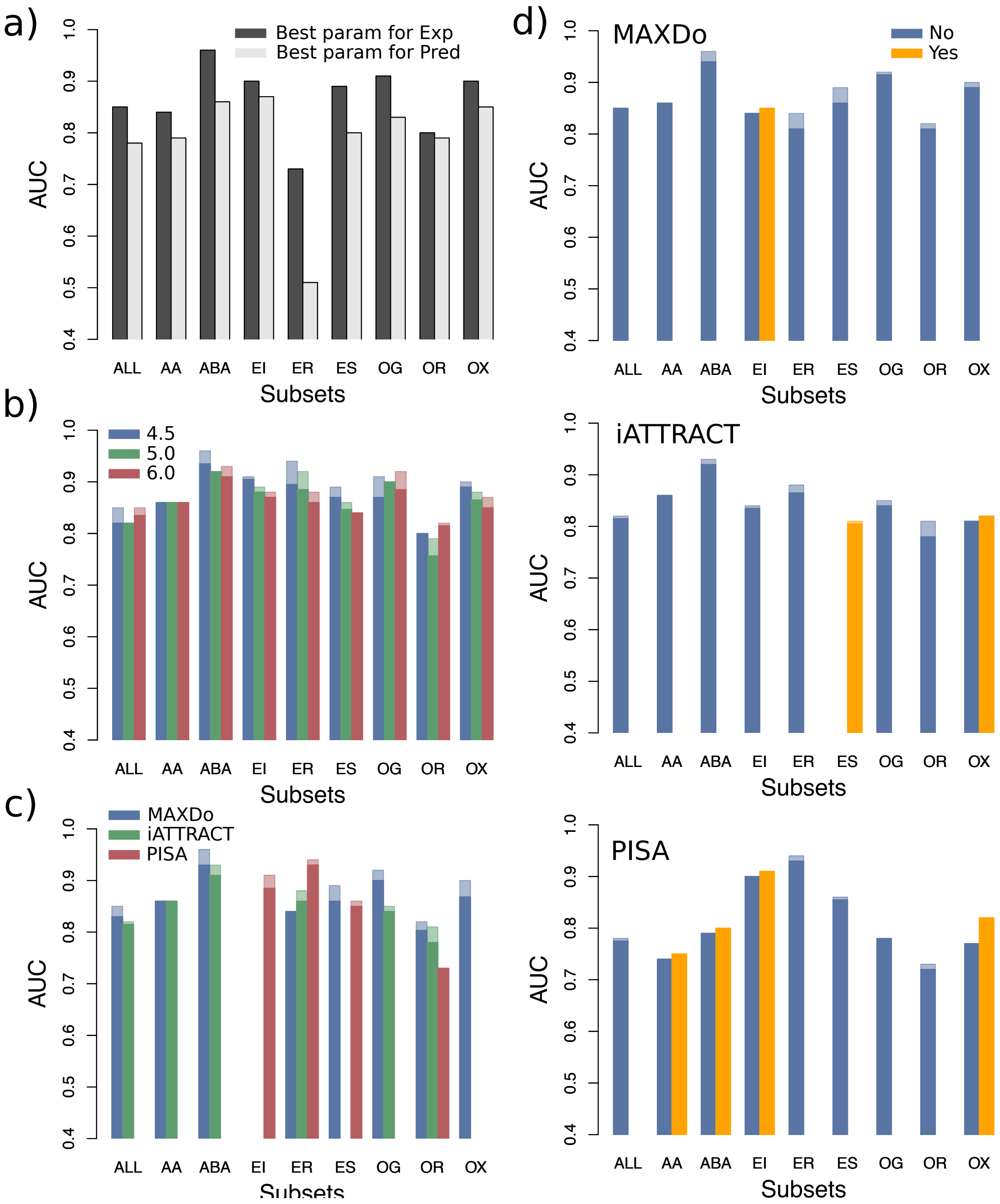


Figure S5: **Detailed predictive performance for PPDBv2, when using the knowledge of the experimental interfaces.** **(a)** Comparison of the AUC values obtained when the parameters were optimized for dealing with experimental interfaces or for dealing with predicted interfaces. The parameters for experimental interfaces are a 6 Å threshold, the MAXDo energy function and no CIPS. They were applied to all classes but EI, where PISA was used instead of MAXDo. The parameters for predicted interfaces are a 5 Å threshold, the MAXDo energy function and CIPS. There are three exceptions: PISA was used for EI, iATTRACT was used for ER and CIPS was not used for OR. **(b-d)** Influence of the individual parameters on the predictive performance. **(b)** Distance threshold used to define docked interfaces. **(c)** Docking energy. **(d)** Presence or absence of the CIPS pair potential, depending of the docking energy. In each plot, for each protein class, we considered the 6 combinations with the highest AUC values. This pool of combinations was divided into 2 to 4 subsets depending on the number of values considered for the parameter. The opaque bars indicate the average AUC values computed over the subsets of combinations. The parts in transparent indicate the maximum values. If a parameter value was not present in the 6 best combinations, then it does not appear on the plot.

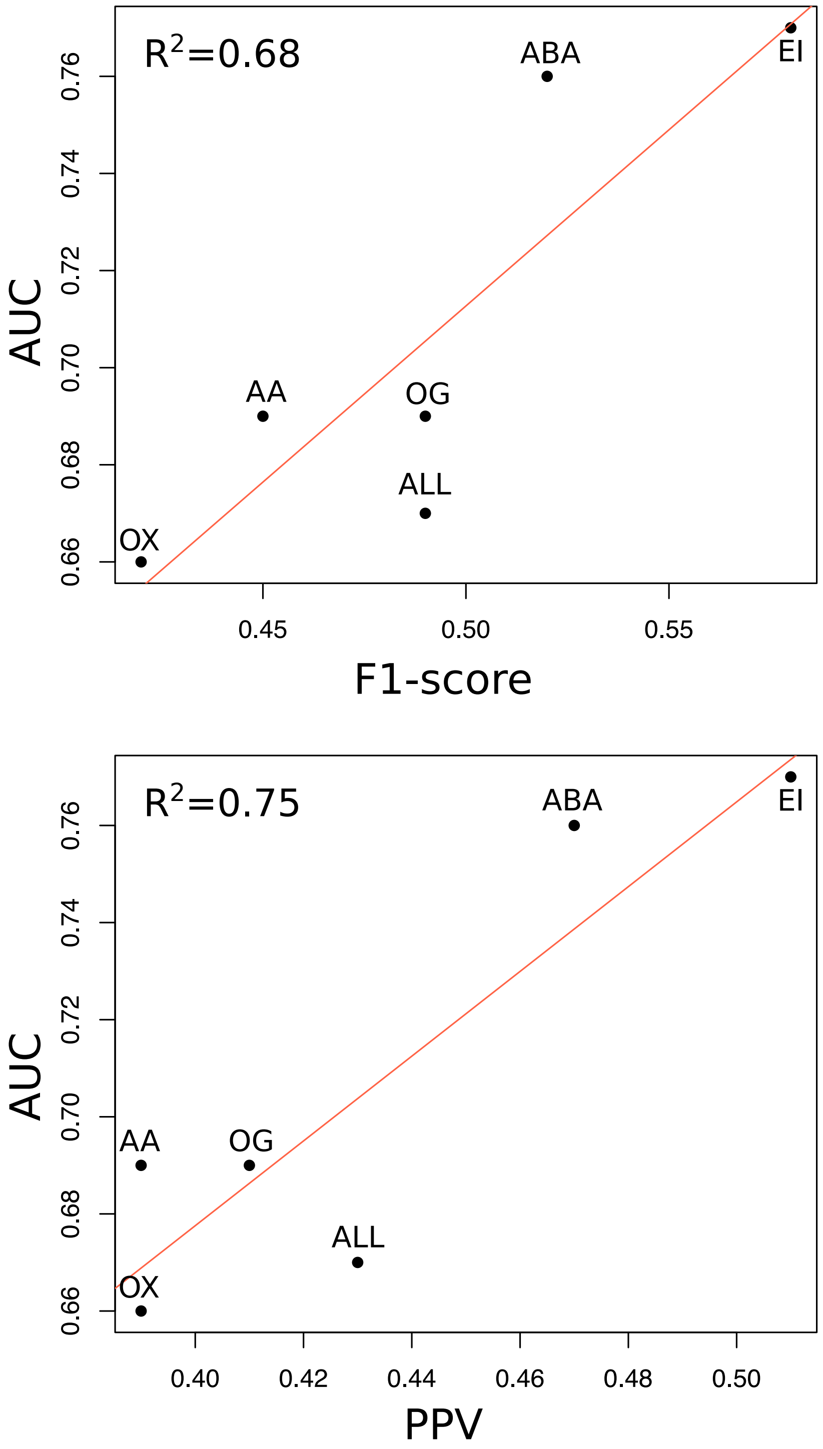


Figure S6: **Influence of the quality of the interface predictions on partner identification** The AUC values are plotted in function of the F1-score and the positive predictive value (PPV) of the predicted RIs, for the whole dataset and a subset of classes (each containing more than 15 proteins). On each plot, the red line corresponds to a linear regression between the two variables, whose adjusted $R^2$ is reported in the top left corner. The scoring strategy is SC-dockSeed-mix and the AUC values correspond to CCD2PI default parameter setting.
